# Estriol is a Stronger Transcriptional Activator than is either 17β-Estradiol or Estrone of Human and Elephant Shark Estrogen Receptor-alpha and Estrogen Receptor-beta transfected into COS-7 Cells

**DOI:** 10.64898/2026.07.03.736429

**Authors:** Ya Ao, Ronn Matthew delos Reyes Cabizares, Michael E. Baker, Yoshinao Katsu

## Abstract

Humans and other vertebrates contain two estrogen receptors (ERs), ERα and ERβ, which mediate the physiological actions of three estrogens: estrone (E1), estradiol (E2) and estriol (E3). Of these three estrogens, *in vivo*, E2 is the strongest transcriptional activator of ERα and ERβ, E1 is next most active, followed by E3. We studied transcriptional activation of human ERα and ERβ by E2, E1 and E3 in African green monkey kidney (COS-7) cells, which we compared with studies of estrogen stimulation of ER transcription in human embryonic kidney (HEK-293) cells. To our surprise, in COS-7 cells, E3 had the lowest half-maximal response (EC50) for human ERα and ERβ than either E2, which was second most active estrogen, or E1. In contrast, for human ERα and ERβ transfected into HEK-293 cells, E2 was the most active estrogen, followed by E1 and E3. Similar results were found in COS-7 cells and HEK-293 cells transfected with elephant shark ERα and ERβ. Thus, under some conditions, E3 is a more active estrogen than either E2 or E1. This suggests that E3 may be a novel physiological ligand for the ER in some mammalian cells, including the placenta, where E3 levels increase substantially during pregnancy.

**Highlights:** The response of the estrogen receptor to its ligand varies according to the specific cell type. In HEK-293 cells, the estrogen receptors demonstrate a high degree of sensitivity to E2, while in COS-7 cells, they exhibit a high degree of sensitivity to E3.

## 1. Introduction

Estrogens are steroid hormones with diverse physiological activities in females and males in humans and other vertebrates [1-5]. Estrogens act by binding to estrogen receptors (ERs)[5,6], which belong to the nuclear receptor family, a large group of transcription factors that includes receptors for progestins, androgens, and corticosteroids [7-9]. To date, two distinct estrogen receptor genes: estrogen receptor alpha (ERα) and estrogen receptor beta (ERβ), have been isolated in mammals and other vertebrates [10-13]. Human ERα and ERβ have strong sequence similarity (97%) in their DNA binding domains, but only about 55% similarity in their ligand-binding domains.

In humans, ERα is expressed in female reproductive tissues (uterus, ovary), breast, kidney and bone, while ERβ is expressed in the male (prostate), ovary, colon, kidney and the immune system [5,14,15]. Interestingly, ERs also are expressed in the brain [15-20]. The response of ERα and ERβ in humans and other vertebrates to 17β-estradiol (E2), the main physiological estrogen, as well as to estrone (E1), which is used as a postmenopausal estrogen in humans, and estriol (E3), which is synthesized by the placenta in primates, (Figure 1) has been studied extensively [5,6,10,21,22].

**Figure 1.**
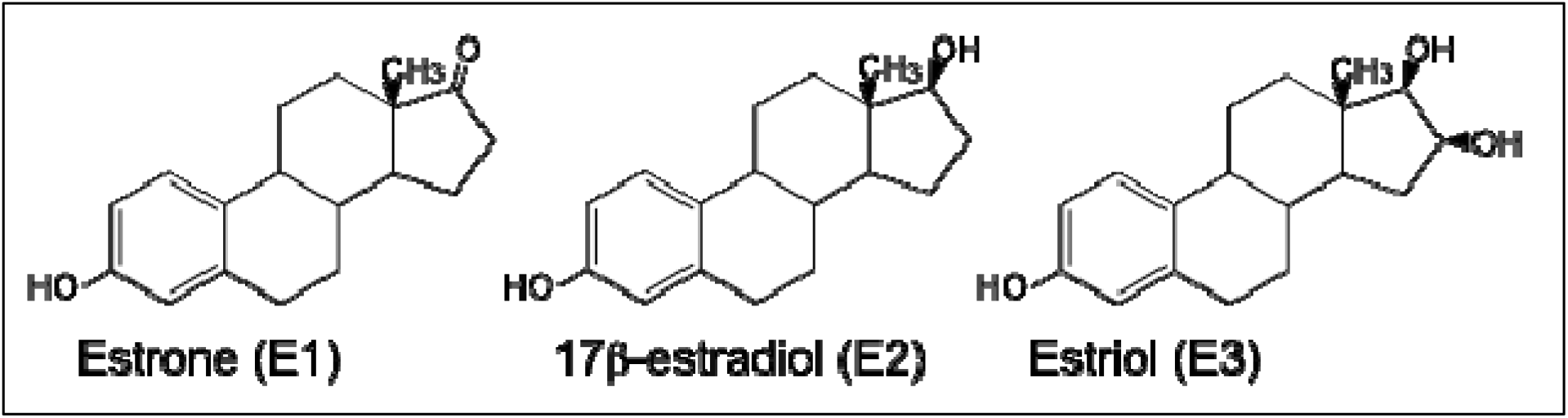
Structures of estrogen. Estrone (E1), 17β-estradiol (E2), and estriol (E3) are the natural estrogens. E1 is a minor female sex hormone and serves mainly as a precursor of E2. E3 is produced in the human placenta, in which E3 is the major estrogen. E2, the major circulating physiological female estrogen, is mainly produced in the ovary [25,26].

ERα and ERβ are activated by binding estrogens, which induce a conformational change in the ER that promotes binding of the hormone-receptor complex to specific DNA sequences leading to the initiation of estrogen-dependent gene transcription. ERα promotes tissue proliferation, while ERβ may act as a suppressor of ERα-mediated proliferation [7,23,24].

To investigate the evolution of estrogen signaling in vertebrates, we compared transcriptional activation in HEK-293 cells of three ERα subtypes ERα1, ERα2, ERα3 and ERβ from elephant shark (*Callorhinchus milii*) with human ERα and ERβ [27]. Due to the low level of transcriptional activation of elephant shark ERs in HEK-293 cells [27], even with high concentrations of E2, we decided to repeat our experiments with *C. milii* ERα and ERβ in monkey kidney (COS-7) cells because COS-7 cells have a high level of responsiveness to exogenously introduced artificial genes [28]. In contrast to our results with HEK-293 cells [27], as reported here, we find that E3 has a lower EC50 for elephant shark ERα and ERβ transfected into COS-7 cells than does either E2 or E1. To determine if this novel activity of E3 is conserved in human ERs, we compared activation of human ERα and ERβ in COS-7 cells and HEK-293 cells by E1, E2 and E3. Here we report that transcriptional activation by either E1, E2, or E3 of human ERα and ERβ transfected into COS-7 cells revealed that E3 has a lower EC50 for human ERα and ERβ than either E2 or E1 has for these ERs. This contrasts with studies with human ERα and ERβ using HEK-293 cells in which E2 has the lowest EC50 for human ERα and ERβ than either E1 or E3 has for these ERs [27]. The low EC50 of E3 compared to E2 for human ERα and ERβ transfected into COS-7 cells suggests that E3 may have previously uncharacterized physiological activities in fetal tissues, which are exposed to amniotic fluid containing high concentrations of E3 [22,29].

## 2. Materials and Methods

### 2.1. Chemical reagents

Estrone (E1), estradiol (E2), and estriol (E3) were purchased from Sigma-Aldrich Corp. (St. Louis, MO). All chemicals were dissolved in dimethylsulfoxide (DMSO). The concentration of DMSO in the culture medium did not exceed 0.1%.

### 2.2. Construction of plasmid vectors

Elephant shark ERs and human ERs have been described previously [27,34]. The full-length estrogen receptor sequences of human (M12674 for ERα, NM_001437 for ERβ), elephant shark (LC068847 for ERα1, XM_007894403 for ERα2, XM_007894404 for ERα3, LC068848 for ERβ), were ligated into the pcDNA3.1 vector (Invitrogen, Carlsbad, CA, USA). The full-coding regions of human ERα (M12674) and ERβ (NM_001437) were amplified by PCR using cDNA synthesized from liver-derived RNA (Takara Bio Inc. Shiga, Japan) as a template. A reporter construct, pGL4.23-4xERE was produced by subcloning of oligonucleotides having 4xERE into the *Kpn*I-*Hind*III site of pGL4.23 vector (Promega Corp. Madison, WI, USA). All cloned DNA sequences were verified by sequencing. Sequencing was performed using a BigDye Terminator Cycle Sequencing kit and analyzed on the Applied Biosystems 3730 DNA Analyzer (Thermo Fisher Scientific Inc. MA, USA).

### 2.3. Empty vector analysis

Analysis of empty vectors for HEK-293 and COS-7 cells were done in triplicate. HEK-293 cells in the presence of 100 nM E1=1.05 ± 0.08, E2=1.02 ± 0.02, E3= 0.86 ± 0.07. COS-7 cells in the presence of 100 nM E1=1.00 ± 0.03, E2=0.98 ± 0.05, E3= 1.04 ± 0.05.

### 2.4. Reporter gene assay and statistical methods

Reporter gene assays were performed in HEK-293 cells [27] or COS-7 cells. Cells were seeded in 24-well plates at 5 × 10^4^ cells/well in phenol-red free Dulbecco’s modified Eagle’s medium with 10% charcoal/dextran-treated fetal bovine serum. After 24 h, the cells were transfected with 400 ng of reporter construct, pGL4.23-4xERE, 25 ng of pRL-TK (as an internal control to normalize the variation in transfection efficiency; contains the *Renilla reniformis* luciferase gene with the herpes simplex virus thymidine kinase promoter, Promega), and 200 ng of pcDNA3.1-receptor using polyethylenimine (PEI) for full-length ERs.

After 5 h of incubation, ligands were applied to the medium at various concentrations. After an additional 43 h, the cells were collected, and the luciferase activity of the cells was measured with the Dual-Luciferase Reporter Assay System. Promoter activity was calculated as firefly (*P. pyralis*)-lucifease activity/sea pansy (*R. reniformis*)-lucifease activity. The values shown are mean ± SEM from three separate experiments, and dose-response data, which were used to calculate the half maximal response (EC50) for each steroid, were analyzed using GraphPad Prism (Graph Pad Software, Inc., San Diego, CA, USA). All experiments were performed in triplicate, independently.

## 3. Results

### 3.1. Transcriptional activation of elephant shark and human ERα1 and ERβ in HEK-293 cells and COS-7 cells

As previously reported [27], the estrogen receptor (ER) from the elephant shark exhibits a high degree of sensitivity to estradiol (E2) in HEK-293 cells, a property that is consistent with estrogen receptors from other species [7,27,35,36,38]. However, low fold-activation values for E2 activation of the ER in HEK-293 cells limited its usefulness for examining the effects of various chemicals. Consequently, an assay utilizing an alter-native cell type was investigated. We decided to use COS-7 cells, which have a higher level of responsiveness to exogenously introduced artificial genes [28,39].

We investigated the response to E1, E2, and E3 of estrogen receptors from elephant sharks and humans transfected into COS-7 cells and found that all three estrogens activated transcription of elephant shark ERs in a dose-dependent manner (Figure 2). Overall, the fold activation values increased for assays in COS-7 cells compared to HEK-293 cells. An unexpected result was the lower EC50 for activation of elephant shark ERs by E3 compared to E1 and E2 (Figure 2, Table 1). The half-maximal response (EC50) for transcriptional activation of elephant shark ERα1 was 1.38 nM for E1, 1.72 nM for E2, and 0.71 nM for E3, whereas the EC50 of elephant shark ERβ was 0.65 nM for E1, 0.63 nM for E2 and 0.28 nM for E3 (Table 1). Elephant shark ERs also showed a higher level of fold activation value with E3 compared to E2 and E1 in COS-7 cells., The lower EC50 value for E3 signifies the higher level of sensitivity to E3 compared to E1 and E2.

**Table 1.** Estrogen activation of full-length shark ERs in HEK293 or COS-7 cells.

|  |  |  | E1 | E2 | E3 |
| --- | --- | --- | --- | --- | --- |
| shark ERα1* | HEK293 | EC50 | 0.49 nM | 0.0059 nM | 0.2 nM |
| | | Fold-Activation (± SEM) | 1.62 (± 0.16) | 1.67 (± 0.04) | 1.57 (± 0.09) |
|  |  | Ratio Efficacy at 10 nM** | 0.98 | **1.00** | 0.95 |
|  |  | Ratio of Potency*** | 0.012 | **1.00** | 0.030 |
| shark ER α2* | HEK293 | EC50 | 0.29 nM | 0.016 nM | 0.31 nM |
| | | Fold-Activation (± SEM) | 1.51 (± 0.12) | 1.79 (± 0.12) | 1.82 (± 0.13) |
|  |  | Ratio Efficacy at 10 nM** | 0.85 | **1.00** | 1.01 |
|  |  | Ratio of Potency*** | 0.055 | **1.00** | 0.052 |
| shark ER α3* | HEK293 | EC50 | 0.25 nM | 0.0093 nM | 0.29 nM |
| | | Fold-Activation (± SEM) | 1.55 (± 0.11) | 1.75 (± 0.08) | 1.87 (± 0.15) |
|  |  | Ratio Efficacy at 10 nM** | 0.88 | **1.00** | 1.06 |
|  |  | Ratio of Potency*** | 0.037 | **1.00** | 0.032 |
| shark ERβ* | HEK293 | EC50 | 0.14 nM | 0.001 nM | 0.02 nM |
| | | Fold-Activation (± SEM) | 2.87 (± 0.16) | 3.23 (± 0.26) | 3.49 (± 0.26) |
|  |  | Ratio Efficacy at 10 nM** | 0.89 | **1.00** | 1.08 |
|  |  | Ratio of Potency*** | 0.007 | **1.00** | 0.05 |
| shark ERα1 | COS-7 | EC50 | 1.38 nM | 1.72 nM | 0.71 nM |
| | | Fold-Activation (± SEM) | 2.66 (± 0.06) | 2.88 (± 0.05) | 4.00 (± 0.14) |
|  |  | Ratio Efficacy at 10 nM** | 0.93 | **1.00** | 1.39 |
|  |  | Ratio of Potency*** | 0.52 | 0.41 | **1.00** |
| shark ERα2 | COS-7 | EC50 | 1.11 nM | 1.58 nM | 0.6 nM |
| | | Fold-Activation (± SEM) | 3.04 (± 0.04) | 3.27 (± 0.16) | 5.08 (± 0.22) |
|  |  | Ratio Efficacy at 10 nM** | 0.93 | **1.00** | 1.55 |
|  |  | Ratio of Potency*** | 0.54 | 0.38 | **1.00** |
| shark ERα3 | COS-7 | EC50 | 1.17 nM | 1.44 nM | 0.51 nM |
| | | Fold-Activation (± SEM) | 2.99 (± 0.08) | 3.24 (± 0.17) | 5.31 (± 0.36) |
|  |  | Ratio Efficacy at 10 nM** | 0.92 | **1.00** | 1.64 |
|  |  | Ratio of Potency*** | 0.44 | 0.35 | **1.00** |
| shark ERβ | COS-7 | EC50 | 0.65 nM | 0.63 nM | 0.28 nM |
| | | Fold-Activation (± SEM) | 2.07 (± 0.15) | 2.1 (± 0.08) | 2.94 (± 0.12) |
|  |  | Ratio Efficacy at 10 nM** | 0.99 | **1.00** | 1.40 |
|  |  | Ratio of Potency*** | 0.43 | 0.44 | **1.00** |

\*Data were from Ao et al. (2026).<sup>26</sup>
\*\*Ratio values were calculated by dividing the fold-activation value of estrogens by the value of 10 nM E2 (in bold).
\*\*\*The potency ratio was calculated using the EC50 values of E2 (HEK293) or E3 (COS-7) set to 1 (in bold).

**Figure 2.**
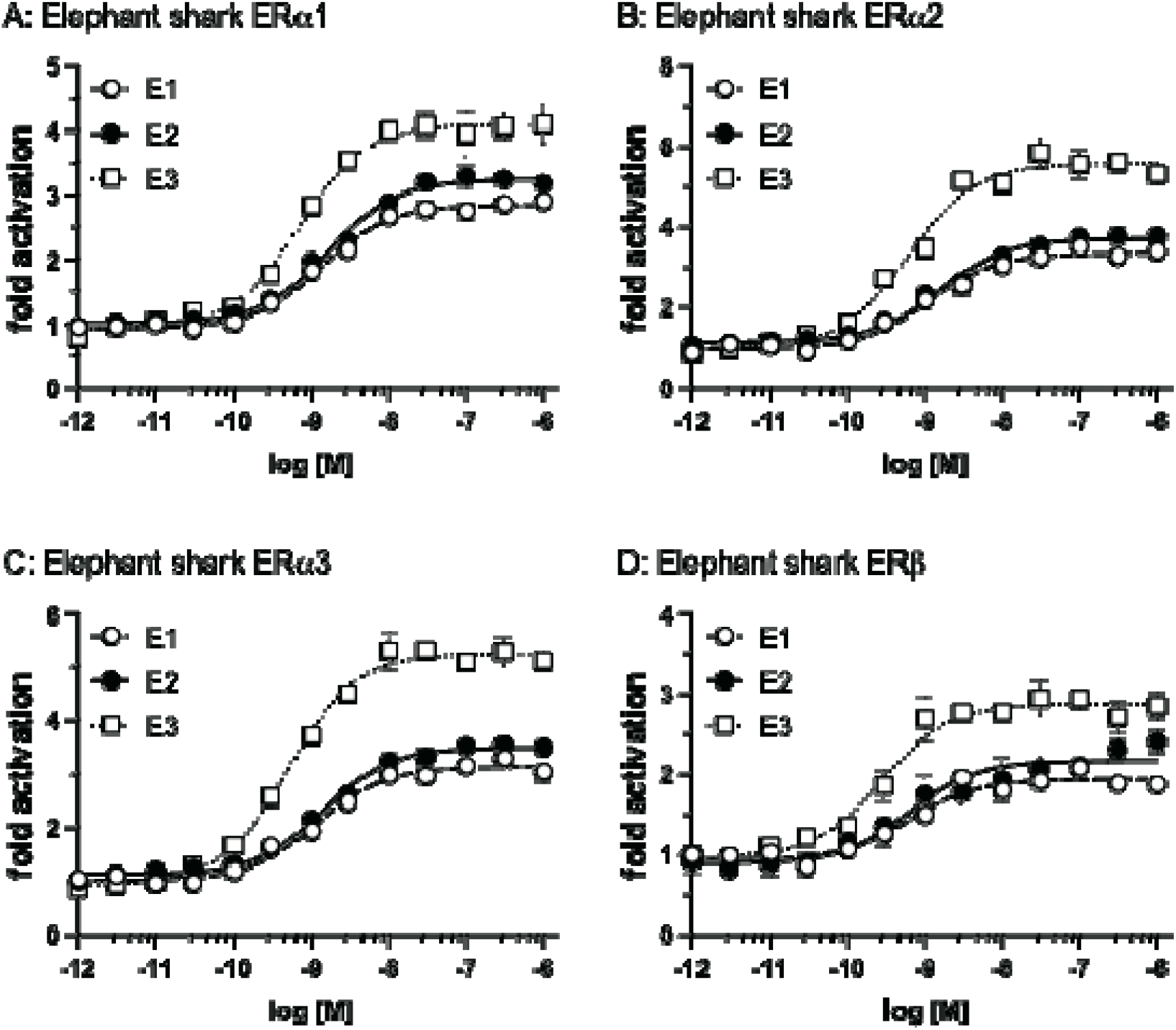
Transcriptional activation of the full-length elephant shark ERs in COS-7 cells. Full-length elephant shark ERs were transfected into COS-7 cells with an ERE-driven reporter gene. Concentration-response profile for ERα1 (A), ERα2 (B), ERα3 (C), and ERβ (B) for E1, E2, and E3 (10^-12^M to 10^-6^M). Data are expressed as a ratio of a vehicle (DMSO) to other test chemicals. Each point represents the mean of triplicate determinations, and vertical bars represent the mean ± SEM.

Next, we investigated estrogen responsiveness of HEK-293 cells and COS-7 cells transfected with human ERα and ERβ. In HEK-293 cells, human ERα and ERβ exhibited a higher sensitivity to E2 compared to E1 and E3 (Figure 3A, B).

**Figure 3.**
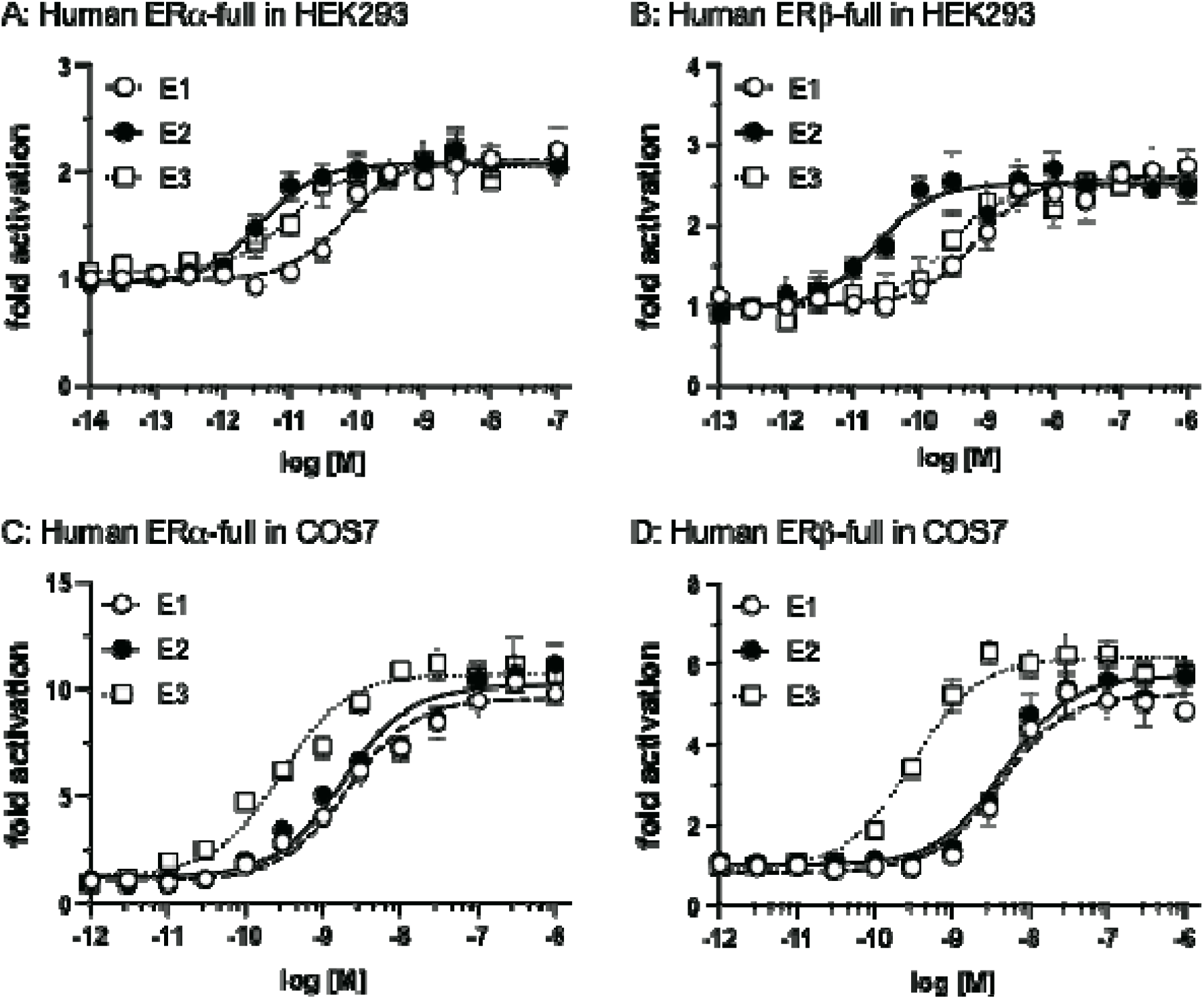
Transcriptional activation of the full-length human ERs in HEK293 or COS-7. Full-length human ERs were transfected into HEK293 or COS-7 cells with an ERE-driven reporter gene. Concentration-response profile for ERα in HEK293 (A), ERβ in HEK293 (B), ERα in COS-7 (C), and ERβ in COS-7 (D) for E1, E2, and E3 (10^-14^ M to 10^-7^ M or 10^-6^ M). Data are expressed as a ratio of a vehicle (DMSO) to other test chemicals. Each point represents the mean of triplicate determinations, and vertical bars represent the mean ± SEM.

The EC50 values for transcriptional activation of human ERα were 0.066 nM for E1, 0.0037 nM for E2, and 0.0095 nM for E3, whereas the EC50 values of human ERβ were 0.7 nM for E1, 0.021 nM for E2, and 0.24 nM for E3 (Table 2). In contrast, both α and β human ERs exhibited a higher degree of sensitivity to E3 compared to E2 and E1, when expressed in COS-7 cells. The EC50 values for transcriptional activation of human ERα were 2.12 nM for E1, 1.84 nM for E2, and 0.29 nM for E3, whereas the EC50 values of human ERβ were 4.25 nM for E1, 4.66 nM for E2, and 0.31 nM for E3 (Figure 3C, D, Table 2).

**Table 2.** Estrogen activation of full-length human ERs in HEK293 or COS-7 cells.

|  |  |  | E1 | E2 | E3 |
| --- | --- | --- | --- | --- | --- |
| human ERα | HEK293 | EC50 | 0.066 nM | 0.0037 nM | 0.0095 nM |
| | | Fold-Activation (± SEM) | 2.1 (± 0.16) | 2.09 (± 0.15) | 1.92 (± 0.05) |
|  |  | Ratio Efficacy at 10 nM* | 1.01 | **1.00** | 0.92 |
|  |  | Ratio of Potency** | 0.056 | **1.00** | 0.39 |
| human ERβ | HEK293 | EC50 | 0.7 nM | 0.021 nM | 0.24 nM |
| | | Fold-Activation (± SEM) | 2.39 (± 0.16) | 2.7 (± 0.21) | 2.2 (± 0.2) |
|  |  | Ratio Efficacy at 10 nM* | 0.89 | **1.00** | 0.81 |
|  |  | Ratio of Potency** | 0.03 | **1.00** | 0.088 |
| human ERα | COS-7 | EC50 | 2.12 nM | 1.84 nM | 0.29 nM |
| | | Fold-Activation (± SEM) | 7.23 (± 0.55) | 7.38 (± 0.49) | 10.86 (± 0.36) |
|  |  | Ratio Efficacy at 10 nM* | 0.98 | **1.00** | 1.47 |
|  |  | Ratio of Potency** | 0.14 | 0.16 | **1.00** |
| human ERβ | COS-7 | EC50 | 4.25 nM | 4.66 nM | 0.31 nM |
| | | Fold-Activation (± SEM) | 4.37 (± 0.31) | 4.72 (± 0.57) | 6.01 (± 0.35) |
|  |  | Ratio Efficacy at 10 nM* | 0.93 | **1.00** | 1.27 |
|  |  | Ratio of Potency** | 0.073 | 0.067 | **1.00** |

\*Ratio values were calculated by dividing the fold-activation value of estrogens by the value of 10 nM E2 (in bold).
\*\*The potency ratio was calculated using the EC50 values of E2 (HEK293) or E3 (COS-7) set to 1 (in bold).

This result for human ERs was similar to that observed in the evaluation of elephant shark ER in COS-7 cells. However, in contrast to the fold activation values observed for elephant shark ER, those of E3 were not significantly higher than those of E1 or E2 for human ERs (Figure 3, Table 2).

## 4. Discussion

The estrogen receptors in Chondrichthyes, cartilaginous fishes with jaws, are basal to lobe-finned fishes and are ancestors of terrestrial vertebrates [40]. Unlike humans, elephant sharks are egg-laying animals. The identification of three estrogen activated ERα subtypes ERα1, ERα2 and ERα3 and ERβ from the sequencing of the elephant shark [40] provided an opportunity to investigate estrogen signaling in a basal jawless vertebrate that evolved about 425 million years ago. For this purpose, we investigated the response to E1, E2, and E3 of ERα1, ERα2, ERα3, and ERβ from elephant shark expressed in HEK-293 cells [27].

These studies revealed that E2 is a stronger than either E1 or E3 in activating transcription of elephant shark estrogen activation of ERα1, ERα2, ERα3, and ERβ, as well as in activating human ERα and ERβ expressed in HEK-293 cells. This is consistent with reports for transcriptional activation of ERs in other vertebrates, and the conserved nature of estrogen signaling pathways [35,41,42]. Indeed, estrogen-dependent transcriptional activity of the ERs from various species, including other sharks, revealed that ERs from many species have a stronger response to E2 than to E3 [37,38,44,45].

In contrast, we find that E3 has a lower EC50 in COS-7 cells compared to its activity in HEK-293 cells in sharks and humans, which diverged over 425 million years ago [40,46]. Our finding that in COS-7 cells, the ER is more sensitive to E3 than either E2 or E1 raises the possibility that E3 may be the biological ligand for the ER in some cells. In amniotic fluid, E3 is the major estrogen in due to its synthesis by the placenta [22,29]. Indeed, E3 levels in amniotic fluid increase dramatically during pregnancy [33], with a peak in the third trimester [47]. Due to the much larger fetal adrenal gland in humans compared to chimpanzees and gorillas, E3 levels in the urine of pregnant women are 4-5 fold higher than levels in the urine of gorillas or chimpanzees [29]. This larger placenta is a characteristic difference between humans and great apes, supporting a role for the placenta and E3 in the divergence of humans and great apes.

E3 is important in the development of mouse brain [30,31]. The higher E3 levels in amniotic fluid in humans compared to either chimpanzees or gorillas may have influenced brain development [32,33], raising the possibility that transcription mediated by E3 contributed to the evolution of human cognition.

## Author Contribution

**Y.A**.: Writing – review & editing, Writing – original draft, Methodology, Investigation, Formal analysis, and Data curation. **R.M.R.C**..: Writing – original draft, Methodology, Investigation, Formal analysis, Data curation. **M.E.B**.: Writing – review & editing, Pro-ject administration, Validation, Supervision, Conceptualization. **Y.K**.: Writing – review & editing, Project administration, Visualization, Validation, Funding acquisition, Supervision, Conceptualization.

## Data availability

All data is freely available on request to Dr. Katsu or to Dr. Baker.

## Acknowledgment

We are indebted to Dr. Lina C. Wang, Minnesota State University, Mankato for a critical reading of this manuscript. We also thank Dr. John Katzenellenbogen for his comments which improved our manuscript. We appreciate the support of this research from colleagues in our laboratory.

